# Cell-Type-Specific Bidirectional Modulation of the Cortico–Thalamo–Cortical Sensory Pathway by Transcranial Focused Ultrasound (tFUS)

**DOI:** 10.64898/2026.03.23.713540

**Authors:** Huan Gao, Sandhya Ramachandran, Mary M. Torregrossa, Bin He

## Abstract

Transcranial focused ultrasound (tFUS) can noninvasively modulate sensory pathways, but the cell-type-specific mechanisms underlying excitatory or inhibitory effects remain unclear. Here, we investigate how tFUS applied to the somatosensory cortex (S1) influences S1 and posterior medial thalamic nucleus (POm) responses to hind paw vibration-tactile stimulation and which neuronal populations mediate these effects. Vibration-tactile stimulation evoked potentials (TEPs) and multi-unit activities (MUA) in S1 and POm were recorded from male rats. Optogenetic tagging was used to identify S1 CaMKII-positive, PV-positive, and SST-positive neurons, while waveform features were used to classify putative excitatory (i.e., regular-spiking units - RSUs) and inhibitory neurons (i.e., fast-spiking units - FSUs) in POm. We found that only S1 CaMKII-positive neurons and POm RSUs responded robustly to tactile stimulation. When tFUS was applied to S1, high pulse repetition frequency (PRF), high duty cycle, and high-pressure stimulation (etFUS) produced excitatory modulation of the sensory pathway, whereas low PRF, low duty cycle, and low-pressure stimulation (itFUS) induced inhibitory effects. Further analyses revealed that excitatory modulation was mediated by activation of S1 CaMKII-positive neurons, while the inhibitory effect arose from their deactivation. These findings demonstrate that tFUS exerts bidirectional, parameter-dependent modulation of a sensory pathway and highlight the critical role of CaMKII-positive neurons in mediating these effects. This study provides mechanistic insight into cell-type-specific neuromodulation by tFUS, particularly in bidirectional modulation of a sensory pathway, and informs the optimization of stimulation parameters for targeted therapeutic interventions.

## Introduction

Transcranial focused ultrasound (tFUS) has emerged as a promising tool for modulating neural circuits [1–7] and developing therapeutic interventions [8–12] by delivering pulsed mechanical energy to targeted brain regions. Compared with other noninvasive neuromodulation techniques, tFUS offers high spatial specificity and the ability to reach deep brain structures.

Previous studies have demonstrated that tFUS can modulate sensory pathways in both animals [13,14] and humans [15,16] depending on stimulation parameters. In human studies, the ability to discriminate tactile vibration frequencies has been shown to improve when excitatory modulation is induced by applying tFUS to the primary somatosensory cortex (S1) [15]. tFUS has also been reported to significantly attenuate the amplitude of somatosensory evoked potentials (SEPs) elicited by median nerve stimulation [16]. In animal studies, targeting specific thalamic nuclei using 1.14 MHz low-intensity focused ultrasound results in suppression of SEPs [14]. In addition, studies in large-animal models have demonstrated that tFUS applied to S1 and its thalamic projections can safely induce transient and reversible suppression of SEPs [13]. These findings suggest that tFUS is capable of modulating sensory processing within the cortico–thalamo–cortical pathway. However, the neuronal mechanisms underlying how different tFUS stimulation parameters produce distinct modulatory effects within sensory pathways remain largely unknown.

Previous studies have suggested that tFUS can produce cell-type-specific effects depending on stimulation parameters. Intracranial recordings in S1 have shown that tFUS can selectively elicit excitatory responses in regular-spiking units (RSUs; putative excitatory neurons) when applied with pulse repetition frequencies (PRFs) above 300 Hz, whereas fast-spiking units (FSUs; putative inhibitory neurons) show no response under these conditions [17]. Activation of RSUs in S1 has also been shown to drive both feedback and feedforward cortico–thalamo–cortical pathways [1]. In addition, studies delivering cell-type-specific doses of tFUS to the cortex have reported activation of several canonical neuronal populations, including Thy1-positive pyramidal excitatory neurons and inhibitory interneurons such as somatostatin-positive (SST) and parvalbumin-positive (PV) neurons [18]. However, whether and how neuronal selectivity is preserved when tFUS modulates sensory processing within the cortico–thalamo–cortical pathway remains unknown.

In this study, vibration-tactile stimulation is applied to the hind paw to activate the sensory pathway. Multi-site intracranial recordings are used to monitor neuronal activity in both S1 and the posterior medial thalamic nucleus (POm) during vibration-tactile stimulation, with or without concurrent tFUS applied to S1 at varying PRFs and duty cycles (DCs) in male rats. Tactile-evoked potentials (TEPs) are analyzed to assess excitatory or inhibitory effects. S1 neurons are classified into CaMKII-positive, PV-positive, and SST-positive populations using optogenetic tagging, while POm neurons are categorized as RSUs or FSUs based on waveform characteristics. Spiking rates from these neuronal populations are measured to evaluate their responses to tactile stimulation and tFUS, providing insight into the mechanisms by which tFUS modulates sensory processing within the cortico–thalamo–cortical (CTC) pathway.

## Methods and Materials

### Experimental Design

#### AAV Virus injection

Male rats (Hsd: WI, Envigo, USA), weighing 250-350g, age 3-6 months, were used as subjects. All animal studies were approved by the Institutional Animal Care and Use Committee at Carnegie Mellon University in accordance with US National Institutes of Health guidelines.

Rats were anesthetized with isoflurane before being mounted on a standard stereotaxic apparatus for viral infusion. Their body temperature was maintained with a heating pad, and heart rate (270-400 bpm) and respiratory rate (50-80 bpm) were monitored during the infusion. The isoflurane was kept at 2.5% with 0.4-0.6 L/min flow rate.

Based on the rat atlas [19], a burr hole was made in the skull over left S1 (AP:-2, ML: 2.5) for each rat. The dura covering the tissue in the hole was carefully removed. PV-ChR2/mCherry (Addgene, No.213945-PHPeB, N = 10), SST-ChR2/mCherry (Addgene, No. 213941-PHPeB, N = 12) or CaMKII-ChrimsonR/mScarlet (Addgene, No. 124651-AAV9, N = 12) were selected to label PV-positive, SST-positive or CaMKII-positive neurons, respectively. 1 μL of the selected virus was injected into the S1 from the left side (DV:1.5mm, angle: 40°) with a speed of 0.2μL/min using a 10 μL microsyringe (Hamilton, No. 7635-01) and a 33-gauge needle (Hamilton, No. 7762-06). After virus infusion, the burr hole was covered by tissue adhesive (3M Vetbond) and the skin was sutured.

#### Surgical Procedures for Anesthetized Recordings

Four weeks following viral infusion, rats underwent surgery to implant electrodes. Fig. 1(A) illustrates the setup for the experiment. Rats were anesthetized with isoflurane before being mounted on the stereotaxic apparatus. Body temperature was maintained with a heating pad and heart rate (270-400 bpm) and respiratory rate (50-70 bpm) were monitored throughout the surgery. Isoflurane was kept at 2% with 0.4-0.6 L/min flow rate during recordings.

**Fig. 1.**
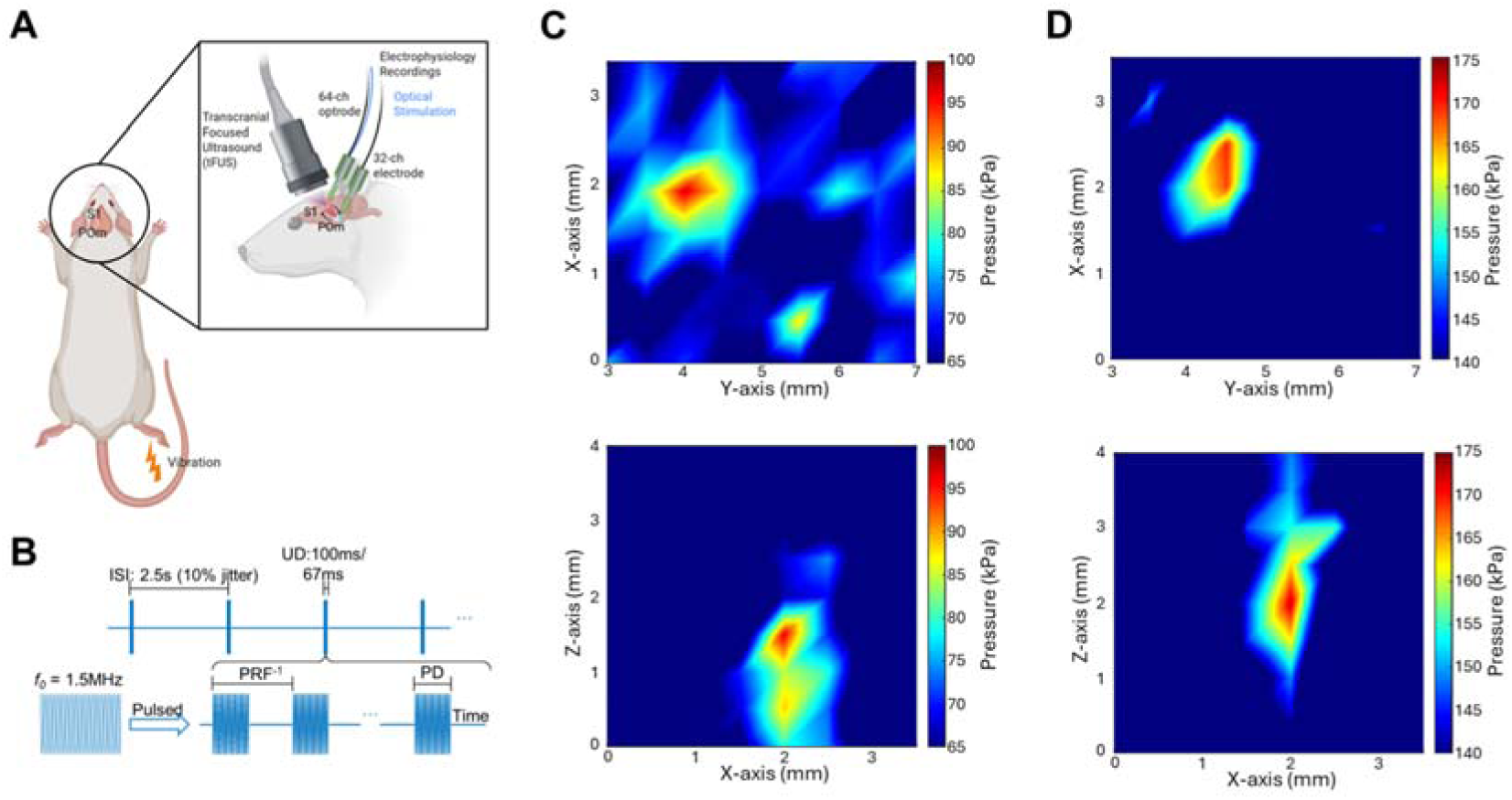
Experimental setup and ultrasound parameter. (A) Experimental setup of intracranial electrodes for recording multi-region signals, optical fiber for activating specific type of neurons and 128-element random array ultrasound transducer. The 64-channel optrode was inserted into left somatosensory cortex (S1) and 32-channel electrode was inserted into left posteromedial of the thalamus (POm). Vibration-tactile stimulation was applied to the right hind paw. (B) Ultrasound parameters used in this study. The inter-sonication interval (ISI) was set at 2.5s with 10% jitter to avoid neuronal adaptation to fixed ISI. Ultrasound duration (UD) was set at 100ms with vibration stimulation or 67ms for tFUS only. Pulse repetitive frequencies (PRFs) and duty cycles (DCs) tested in this study were 30Hz with 0.6%DC and 3000Hz with 60% DC. Pulse durations (PD) were calculated by PRF*DCs. (C-D) Ex-vivo hydrophone peak-to-peak pressure amplitude z-axis (upper) and y-axis (bottom) scan at the estimated targeted brain region with low pressure (C, ∼98kPa) and high pressure (D, ∼163kPa).

To explore the neuronal responses to tFUS, neuronal signals from the S1 and POm were recorded. The POm is a higher-order thalamic area that projects to the cortex and receives driver input from the cortex [21]. The S1 burr hole was remade exactly at the virus injection site (AP: −2, ML: 2.5), and the POm hole was made at AP: −4.5 and ML: 2.6 in the skull for each rat. The dura covering the holes was carefully removed. A 64-channel Optrode (A1×64-Edge-6mm-22.5-177-OXA64LP, Neuronexus, Ann Arbor, MI, USA) was inserted into S1 from the left side (angle: 40°) to make sure the signals were recorded from the virus injection site. The 32-channel NeuroNexus microelectrode (A1×32-Poly3-10mm-50-177, NeuroNexus) was inserted into POm from back side (angle: 28°, DV:5.8) to leave space for the transducer. The transducer was set vertically over the skull after the insertion of electrodes.

#### Vibration-Tactile Stimulation

To test the tFUS effect on the sensory CTC pathway, vibration-tactile stimulation was used as it has been widely applied to study the functional network integrity of ascending sensory pathways [21–23]. The vibration motor was placed under the right hind paw. A waveform generator was used to drive the motor producing vibration-tactile stimulation. The output was set at 5V with frequency of 100Hz or 500Hz. The duration of stimulation was 100ms.

#### Transcranial focused ultrasound stimulation (tFUS)

The customized 128-element random array ultrasound transducer H276 (*f*_0_: 1.5 MHz, −3dB axial specificity: 1.36 mm and lateral specificity: 0.46 mm, manufactured by Sonic Concepts, Inc., Bothell, WA, USA) was used. The transducer was driven by a Vantage 128 research ultrasound platform (Verasonics, Kirkland, WA, USA) using a DL-260 connector to steer the ultrasound beam. Ex-vivo scanning was performed using the customized hydrophone-based 3D ultrasound pressure mapping system (HNR500, Onda Corporation, Sunnyvale, CA, USA). To mimic the *in-vivo* experimental setup, a rat skull sample was placed in degassed water, with the transducer placed vertically over the skull and the hydrophone measuring the ultrasound pressure beneath the skull. The peak-to-peak in-situ ultrasound pressure magnitude with beam steering was 98 kPa for the low-pressure level (Fig. 1C) and 163kPa for the high-pressure level (Fig. 1D) and spatial-peak temporal average (I_SPTA_) was 32.76 mW/cm^2^ and 54.49mW/cm^2^ respectively with 1500 kHz PRF, 30% DC and 67ms UD.

In our study, the inter-sonication interval (ISI) was set at 2.5s for each trial with 10% jitter to avoid entraining any response, and the UD was kept at 100 ms for conditions with tactile stimulation and 67ms for tFUS only condition (Fig. 1B). 3000Hz PRF with 60% DC and 30Hz PRF with 0.6% DC were used with either low pressure or high pressure. Each recording contained 250 trials for tFUS with tactile stimulation or 500 trials for tFUS only. The order of conditions was randomized.

### Data acquisition and preprocessing

All data were recorded using a commercial multi-channel neural signal acquisition system (Tucker-Davis Technologies, Alachua, FL, USA). For evoked potential analysis, recorded local field potentials (LFP) were bandpass filtered between 3Hz and 300Hz with a zero-phase infinite impulse response filter then notch filtered with 60Hz. For spike analysis, recorded MUAs were bandpass filtered between 300 Hz and 6 kHz. Spike sorting was performed using Offline Sorter (Plexon). Spike waveforms were detected beyond the threshold of −3.5 standard deviations from the Mean of Peak Heights Histogram, and clustering was performed using k-means. The spike waveforms and timestamps were stored for further analysis in MATLAB (R2022a, MathWorks, USA).

### Signal Processing

#### Tactile Evoked Potentials (TEPs)

Tactile-evoked potentials (TEPs) were analyzed to evaluate neural responses to vibration-tactile stimulation with the epoch from 100 ms before the trial onset time to 500 ms after. The filtered LFPs were then normalized using a z-score transformation relative to a baseline period of 100 ms prior to the onset of tFUS stimulation for each trial. The z-scored signals were subsequently averaged across recording channels and across subjects to obtain the mean TEP waveform for each experimental condition.

#### Neuronal type classification and response quantification

Only neurons that fired more than 800 spikes per recording session were included for further clustering analysis. To assess neuronal responses to vibration-tactile stimulation and tFUS, neuronal types were first identified. For neurons recorded in S1, optical stimulation was applied in opto-tagged rats for cell-type identification. Opto-tagged neurons were identified by a significant increase in firing rate during optical stimulation compared with the 100 ms pre-stimulation baseline (p < 0.05) and a maximum spike latency of 8 ms following stimulation onset [24]. Optical stimulation was delivered using red light (630 nm) for CaMKII-positive neurons or blue light (465 nm) for SST-positive or PV-positive neurons (50 ms duration; PlexBright LED Module, Plexon, Dallas, TX, USA). Optical stimulation was conducted immediately prior to the tFUS session. Sorted neurons were identified as the same neuron types identified by opto-tagging when their waveforms during the tFUS session showed high similarity to those of previously identified opto-tagged neurons (correlation coefficient > 0.9)[25].

For the identification of neurons in POm, spike waveform features were analyzed. Specifically, the initial phase (IP, from spike onset to the first re-crossing of the baseline) and the afterhyperpolarization period (AHP, the interval from the end of the IP to its subsequent re-crossing of the baseline) were measured. A k-means clustering algorithm was then applied to classify neuronal types based on spike waveform characteristics, as inhibitory neurons typically exhibit shorter spike durations and narrower waveforms. Based on this classification, neurons were categorized as regular-spiking units (RSUs) or fast-spiking units (FSUs).

Neuronal firing rates were calculated, and peri-stimulus time histograms (PSTHs; bin size: 25 ms) were generated to quantify neuronal responses to tactile stimulation and tFUS.

### Statistical analysis

All the results showed in figures with statistics are mean ± S.E.M. The TEPs were compared using the Kruskal-Wallis test and post hoc pairwise comparisons were performed using the Wilcoxon signed-rank test. The spiking rates were normalized to the pre-stim period (100 ms pre-stim) for comparison across different conditions using the Kruskal-Wallis H test. *Post hoc* one-tail two-sample Wilcoxon tests with the Bonferroni correction were employed. Additionally, normalized spike rates within defined response windows were compared using one-way ANOVA with Bonferroni post hoc correction. The statistical significance level was set at 0.05.

## Results

### Tactile stimulation evoked potentials (TEPs) and neuronal responses

Vibration-tactile stimulation reliably evoked responses in both S1 and POm, as evidenced by clear tactile-evoked potentials (TEPs; Fig. 2A, N = 34 rats). In S1, the first negative peak occurred at approximately 22 ms (N22) after stimulation onset in both the 100 Hz (v100) and 500 Hz (v500) vibration groups, and both were significantly different from baseline (p < 0.05), with no significant difference observed between the two frequencies. TEPs in POm showed a clear negative peak around 22 ms only in the 100 Hz group. Although a negative peak was observed in the 500 Hz group, it did not differ significantly from baseline. Given the greater sensitivity of POm responses to 100 Hz stimulation, subsequent experiments with tFUS focused primarily on this frequency.

**Fig. 2.**
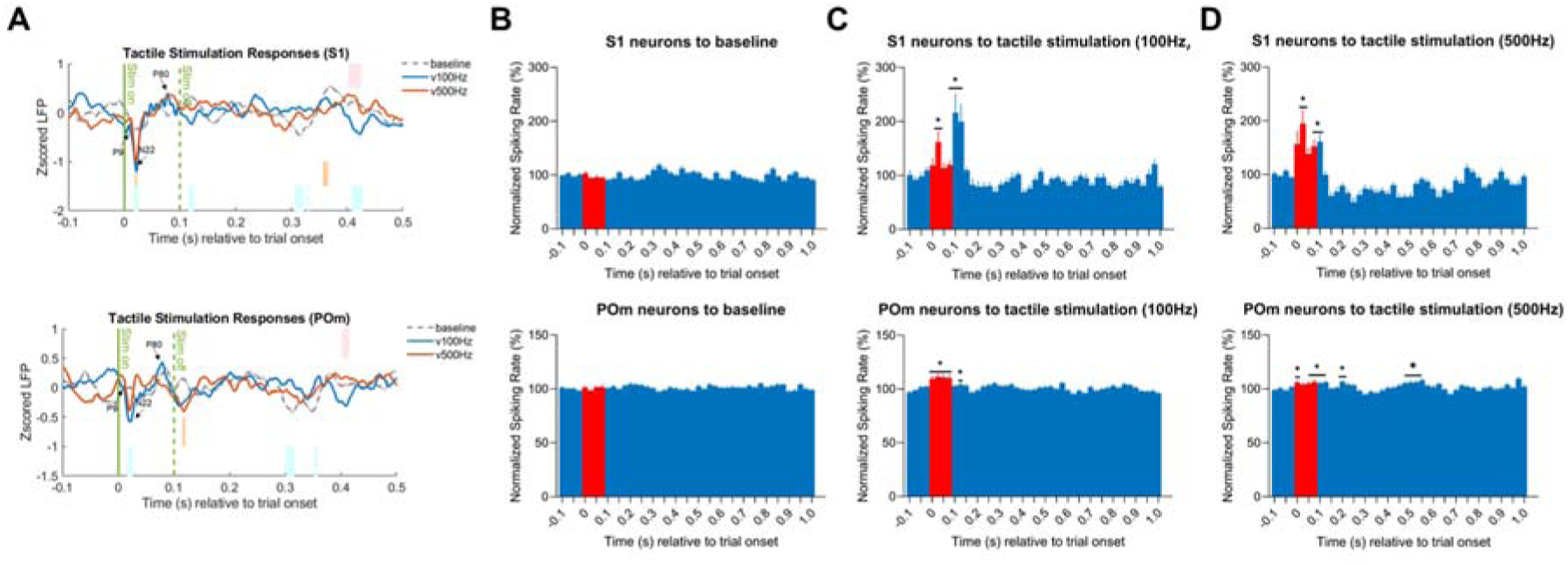
Vibration-tactile stimulation evoked potentials (TEPs) and neuronal responses. (A) TEPs in S1 and POm to baseline (grey, no tactile stimulation), tactile stimulation with 100Hz (blue) and 500Hz (orange) frequencies. The stimulation duration is 100ms. The blue, orange and pink bars indicate significant difference between baseline vs v100Hz, baseline vs v500Hz and v100Hz vs v500Hz, respectively (p<0.05). (B-D) S1 (upper) and POm (bottom) neuronal responses to baseline (B), tactile stimulation with 100Hz (C) and 500Hz (D) frequencies. The * indicates a significant difference in the normalized firing rate of the bin/bins compared to the 100 ms pre-stimulation baseline firing rate (p < 0.05). Bin size is 25 ms. Red bars indicate bins during the stimulation period.

Analysis of multi-unit activity (MUA; Fig. 2B–D) revealed that neurons in both S1 (v100: n = 56; v500: n = 54 from 34 rats) and POm (v100: n = 124; v500: n = 107 from 34 rats) responded significantly to 100 Hz and 500 Hz vibration stimulation compared with their respective baseline groups (S1: n = 101; POm: n = 375 from 34 rats). Notably, POm neurons exhibited responses approximately 25 ms earlier than S1 neurons, consistent with the ascending sensory pathway from thalamus to cortex.

Following neuronal type identification, the responses of specific neuron types were analyzed (Fig. 3). CaMKII-positive (n = 24 from 12 rats), SST-positive (n = 15 from 12 rats), and PV-positive (n = 10 from 10 rats) neurons in S1 were identified based on their responses to optical stimulation (Fig. 3A–C, Fig. S1). Tactile-evoked responses showed distinct patterns across these cell types (Fig. 3D–F), with only CaMKII-positive neurons exhibiting significant responses to vibration-tactile stimulation, while SST-positive and PV-positive neurons showed no detectable responses. In POm, RSUs and FSUs were classified based on their IP and AHP waveform features (Fig. 3G). Only RSUs (n = 63 from 34 rats) displayed robust tactile-evoked responses, while FSUs (n = 61 from 34 rats) remained unresponsive.

**Fig. 3.**
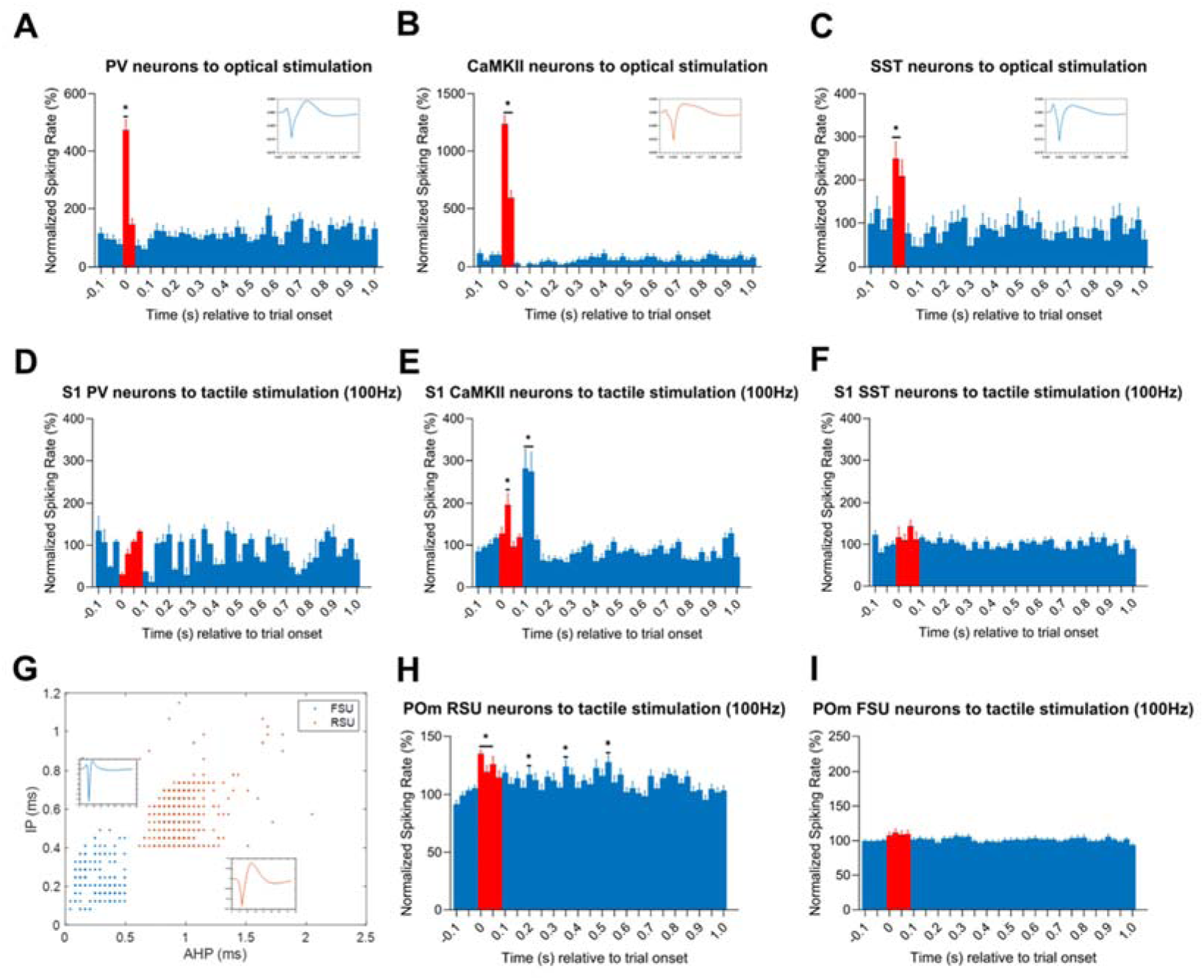
Identification of neuronal subtypes and their responses to tactile stimulation. (A-C) S1 neuronal subtype identification using optical stimulation in opto-tagged rats. The PV-positive (A), CaMKII-positive (B), and SST-positive (C) neurons exhibited more than twofold increase in firing rate relative to the pre-stimulation period. Insets display the representative spike waveforms of the recorded neurons. (D-E) Responses of identified S1 PV-positive (D), CaMKII-positive (E), and SST-positive (F) neurons to 100Hz vibration-tactile stimulation. (G) POm neuronal subtype identification based on waveform characteristics. RSU and FSU clusters based on IP and AHP calculation of waveforms using k-means. The * indicates a significant difference in the normalized firing rate of the bin/bins compared to the 100 ms pre-stimulation baseline firing rate (p < 0.05). Bin size is 25 ms. Red bars indicate bins during the stimulation period.

### TEPs and neuronal responses to tFUS with High PRF and DC

The TEPs and neuron type-specific responses to tFUS with a 3 kHz PRF and 60% duty cycle (DC) were analyzed (Fig. 4) to examine the modulatory effects of tFUS. Both high- and low-pressure levels were tested in combination with tactile stimulation to evaluate how tFUS intensity influences cortical and thalamic responses. Based on the TEPs, both S1 and POm exhibited a significant increase in the responsive peak (N22) in the high-pressure tFUS plus tactile stimulation group (v100+H3kHz) compared with tactile stimulation alone (Fig. 4A). In S1, the N22 peak was also significantly larger than in the tactile stimulation combined with low-pressure tFUS group (v100+L3kHz), indicating a pressure-dependent modulatory effect of tFUS.

**Fig. 4.**
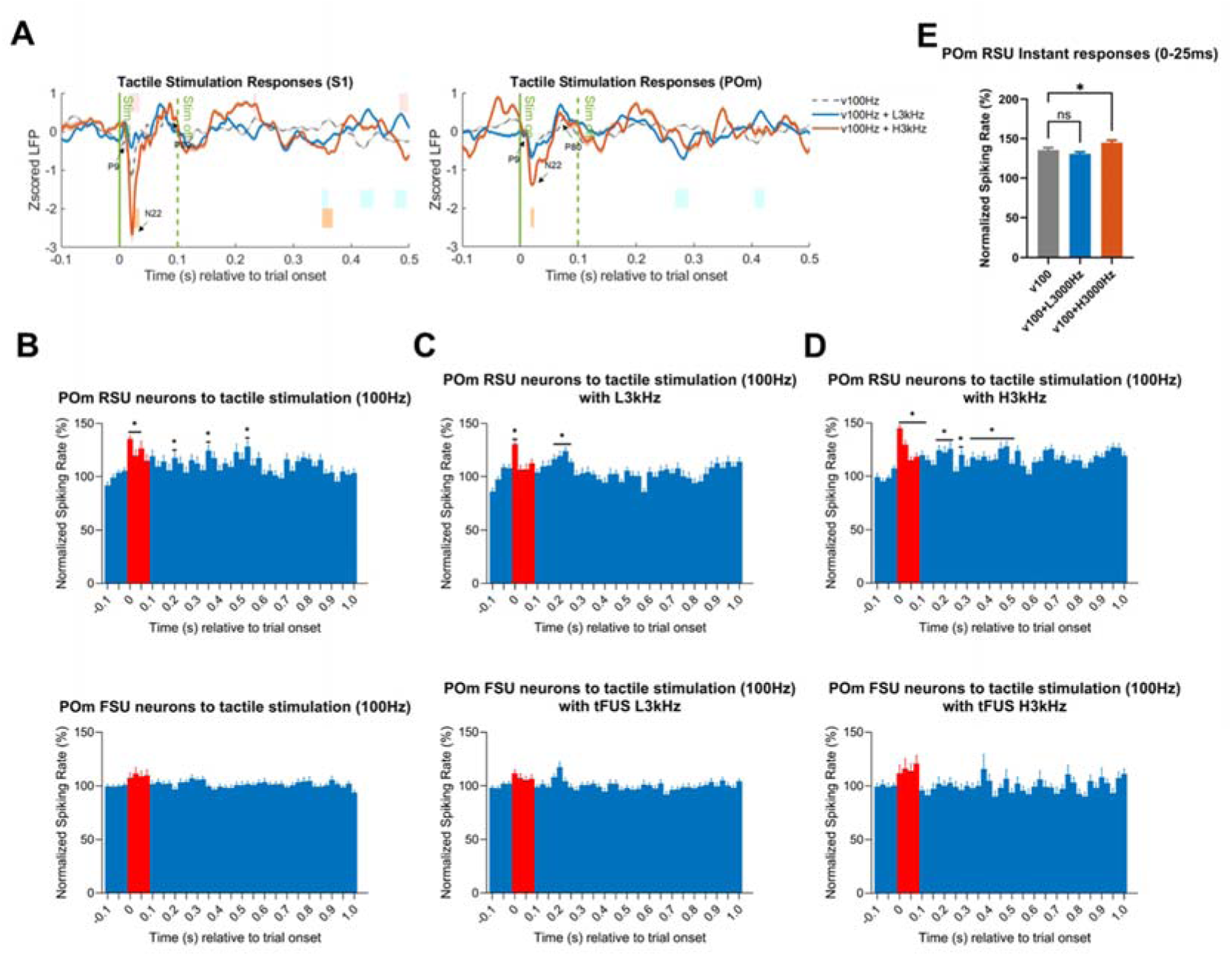
TEPs and neuronal responses to 100 Hz vibration-tactile stimulation combined with tFUS (3 kHz PRF, 60% duty cycle). (A) TEPs recorded in S1 and POm during 100 Hz vibration-tactile stimulation alone (v100Hz, gray), 100 Hz stimulation combined with low-pressure tFUS (v100Hz+L3kHz, blue), and 100 Hz stimulation combined with high-pressure tFUS (v100Hz+H3kHz, orange). Stimulation duration was 100 ms. Blue, orange, and pink bars indicate significant differences between v100Hz vs. v100Hz+L3kHz, v100Hz vs. v100Hz+H3kHz, and v100Hz+L3kHz vs. v100Hz+H3kHz, respectively (p < 0.05). (B-D) POm RSU (upper) and FSU (bottom) neuronal responses to v100Hz only (B), v100Hz+L3kHz(C) and v100Hz+H3kHz (D). The * indicates a significant difference in normalized firing rate of the bin(s) compared to the 100 ms pre-stimulation baseline (p < 0.05). Bin size was 25 ms. Red bars denote bins during the stimulation period. (E) Comparison of normalized RSU firing rate in the first 25 ms window among v100Hz, v100Hz+L3kHz, and v100Hz+H3kHz conditions. * indicates significant difference between v100 and v100+H3kHz groups (p<0.05).

As ultrasound pressures above 130 kPa can induce displacement of intracranial electrodes, which can be detected in the recorded signals [1,26], we focused our analysis on POm neuronal responses to tactile stimulation combined with tFUS. Consistent with previous observations, only POm RSUs showed significant responses to tactile stimulation with or without tFUS, whereas FSUs exhibited no detectable responses (Fig. 4B–D). To further quantify the effect, normalized firing rates within the 0–25 ms window following tactile stimulation and tFUS onset were compared across groups. High-pressure tFUS elicited significantly higher responses than the other groups (Fig. 4E), indicating that tFUS with high PRF, high DC, and high pressure produces an excitatory modulatory effect on the sensory pathway.

### TEPs and neuronal responses to tFUS with Low PRF and DC

The TEPs and neuron type-specific responses to tFUS with a 30 Hz PRF and 0.6% duty cycle (DC) were analyzed (Fig. 5) to examine modulatory effects at both high- and low-pressure levels. Based on the TEPs, both S1 and POm exhibited a significant decrease in the responsive peak (N22) in the low-pressure tFUS plus tactile stimulation group (v100+L30Hz) compared with tactile stimulation alone and tactile stimulation combined with high-pressure tFUS (v100+H30Hz) (Fig. 5A).

**Fig. 5.**
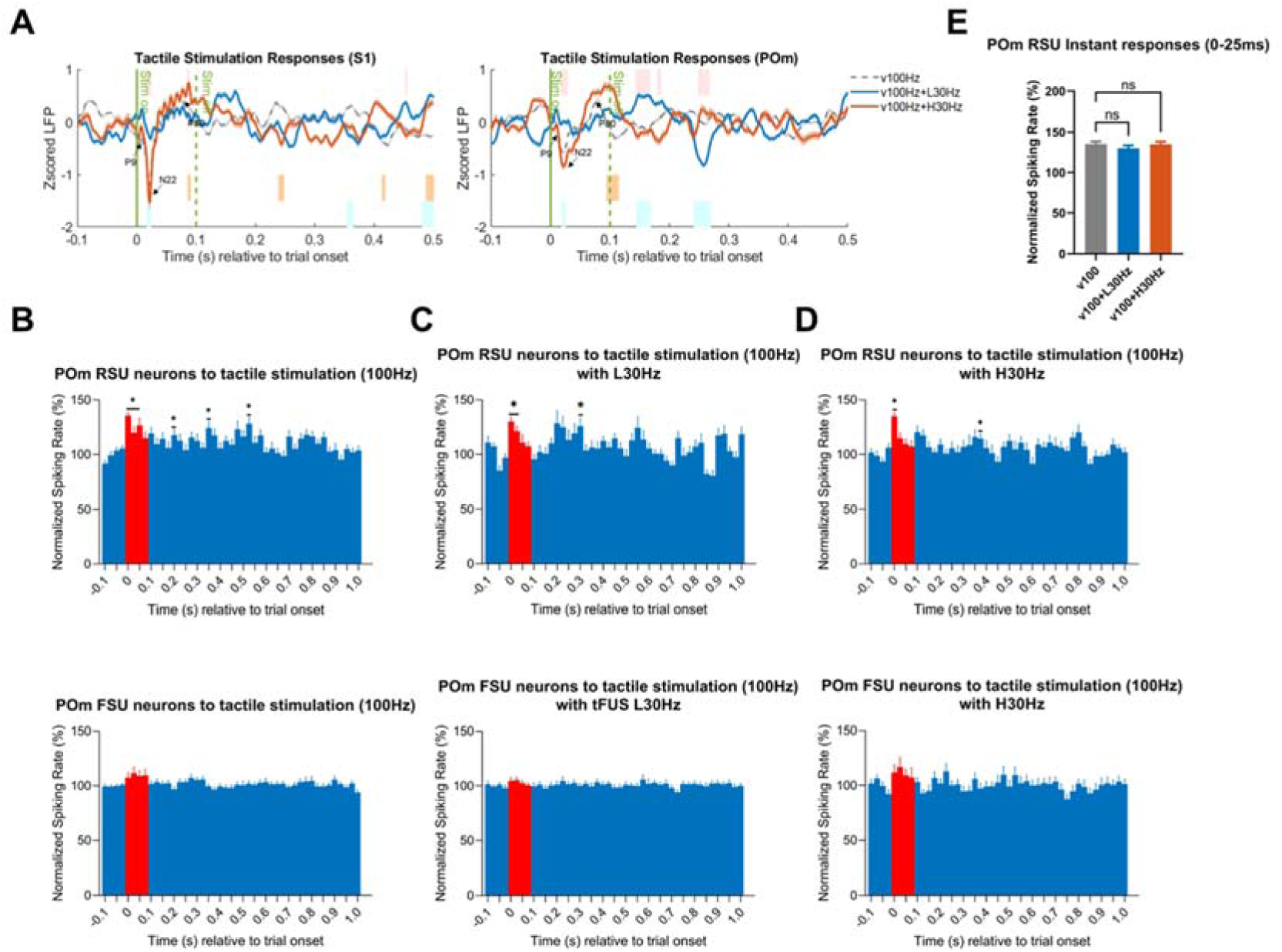
TEPs and neuronal responses to 100 Hz vibration-tactile stimulation combined with tFUS (30 Hz PRF, 0.6% duty cycle). (A) TEPs recorded in S1 and POm during 100 Hz vibration-tactile stimulation alone (v100Hz, gray), 100 Hz stimulation combined with low-pressure tFUS (v100Hz+L30Hz, blue), and 100 Hz stimulation combined with high-pressure tFUS (v100Hz+H30Hz, orange). Stimulation duration was 100 ms. Blue, orange, and pink bars indicate significant differences between v100Hz vs. v100Hz+L30Hz, v100Hz vs. v100Hz+H30Hz, and v100Hz+L30Hz vs. v100Hz+H30Hz, respectively (p < 0.05). (B-D) POm RSU (upper) and FSU (bottom) neuronal responses to v100Hz only (B), v100Hz+L30Hz(C) and v100Hz+H30Hz (D). The * indicates a significant difference in normalized firing rate of the bin(s) compared to the 100 ms pre-stimulation baseline (p < 0.05). Bin size was 25 ms. Red bars denote bins during the stimulation period. (E) Comparison of normalized RSU firing rate in the first 25 ms window among v100Hz, v100Hz+L30Hz, and v100Hz+H30Hz conditions. No significant difference was observed among groups (p>0.05).

Consistently, only POm RSUs responded to tactile stimulation with or without tFUS (Fig. 5B–D), and there was no significant difference in normalized firing rates within the 0–25 ms window following tactile stimulation and tFUS onset across groups (Fig. 5E).

Overall, these results indicated that tFUS with low PRF, low duty cycle, and low pressure exerted an inhibitory modulatory effect on the sensory pathway.

### Cell-type-specific mechanisms underlying bidirectional modulation of the sensory pathway by tFUS

We found that tFUS modulates the sensory pathway in a parameter-dependent manner, producing excitatory effects with high PRF, high DC, and high pressure, and inhibitory effects with low PRF, low DC, and low pressure, raising the question of which neurons are responsible for these differential effects.

The excitatory effects could not be directly assessed in S1 neurons due to stimulation-induced vibration [1,26]. Therefore, we analyzed POm neuronal responses while selectively activating CaMKII-positive, PV-positive, and SST-positive neurons in S1 to investigate the cell-type-specific effects of tFUS (Fig. 6). POm RSUs responded to tactile stimulation both with and without optical activation of CaMKII-positive (n = 24 from 12 rats), PV-positive (n = 20 from 10 rats), or SST-positive (n = 17 from 12 rats) neurons in S1, whereas POm FSUs showed no responses (CaMKII-positive, n = 23 from 12 rats; PV-positive, n = 20 from 10 rats; SST-positive, n = 25 from 12 rats) (Fig. 6A–D). Additionally, optical activation of S1 CaMKII-positive neurons induced a significant increase in firing of POm RSUs within the 0–25 ms window following tactile stimulation onset (Fig. 6E). These results provide indirect evidence that the excitatory modulation of the sensory pathway by tFUS is mediated through activation of CaMKII-positive neurons in S1.

**Fig. 6.**
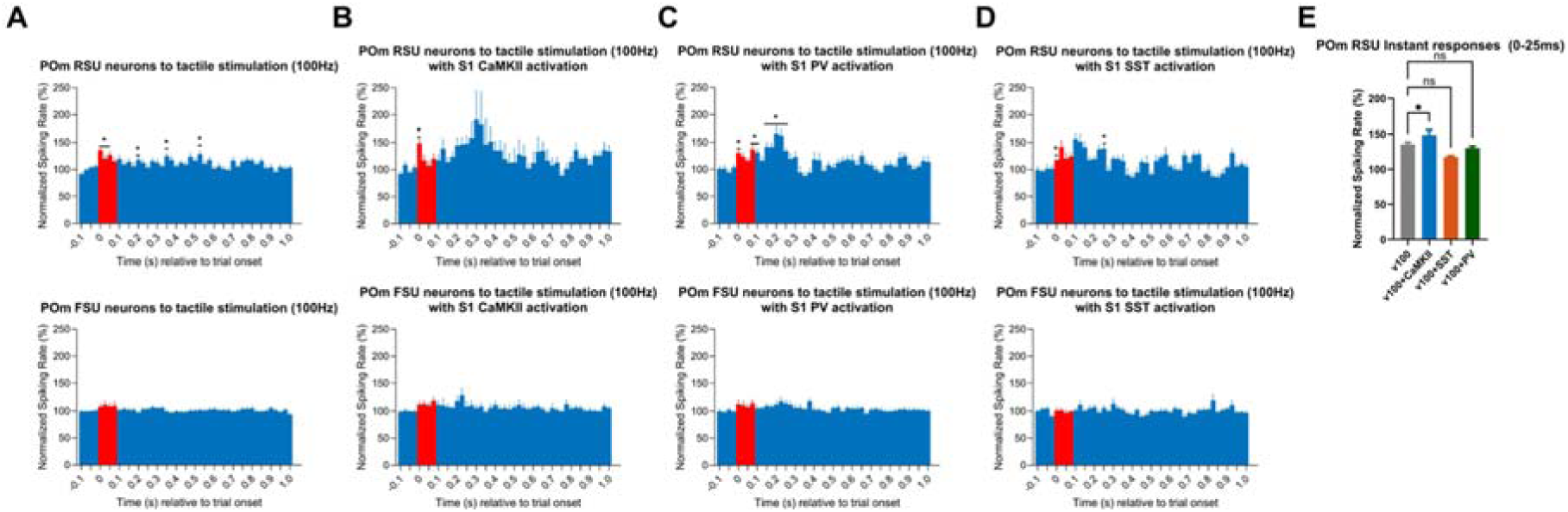
POm RSU and FSU neuronal responses to 100 Hz vibration-tactile stimulation combined with selective activation of S1 neurons via optical stimulation in opto-tagged rats. (A-D) POm RSU (upper) and FSU (bottom) neuronal responses to v100Hz only (A), v100Hz with S1 CaMKII-positive neuronal activation (B), v100Hz with S1 PV-positive neuronal activation (C) and v100Hz with S1 SST-positive neuronal activation (D). The * indicates a significant difference in normalized firing rate of the bin(s) compared to the 100 ms pre-stimulation baseline (p < 0.05). Bin size was 25 ms. Red bars denote bins during the stimulation period. (E) Comparison of normalized RSU firing rate in the first 25 ms window among v100Hz, v100Hz+S1 CaMKII-positive activation, v100Hz+ S1 SST-positive activation and v100Hz+ S1 PV-positive activation conditions. * indicates significant difference between v100 and v100Hz+S1 CaMKII-positive activation groups (p<0.05).

POm neuronal responses did not reveal the mechanism underlying the inhibitory effect. Therefore, we directly analyzed S1 neuron-type-specific responses to low-pressure tFUS to investigate this effect (Fig. 7). S1 CaMKII-positive neurons (n = 20 from 12 rats) exhibited a decrease in firing rate approximately 25 ms after tFUS onset, whereas SST-positive (n = 15 from 12 rats) and PV-positive neurons (n = 14 from 10 rats) showed no responses. To further determine which tFUS parameters were critical for this inhibitory effect, CaMKII-positive responses were examined under high PRF with high DC (n = 25 from 12 rats), high PRF with low DC (n = 22 from 12 rats) and low PRF with high DC (n = 19 from 12 rats) at low pressure (Fig. S2). No decrease in firing rate was observed under these conditions, indicating that the inhibitory effect is specifically induced by the combination of low PRF, low DC, and low pressure.

**Fig. 7.**
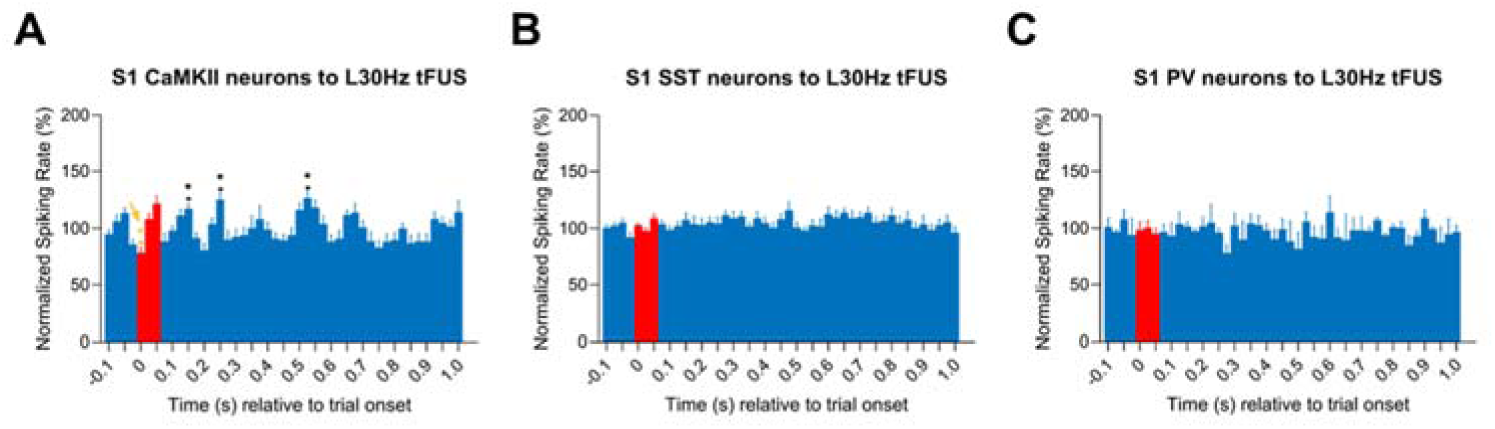
S1 neuronal responses to low-pressure tFUS (30 Hz PRF, 0.6% duty cycle). (A–C) Responses of S1 CaMKII-positive (A), SST-positive (B), and PV-positive (C) neurons to tFUS. Yellow *(pointed by yellow arrow) indicates a significant decrease in normalized firing rate compared to the 100 ms pre-stimulation baseline, whereas black * indicates significant increases relative to baseline (p < 0.05). Bin size was 25 ms. Red bars denote bin windows during the stimulation period.

## Discussion

In this study, we recorded neuronal activity in both S1 and POm during vibration-tactile stimulation, with and without tFUS. We observed that only S1 CaMKII-positive neurons and POm RSUs responded robustly to tactile stimulation. We then examined neuronal responses to high- and low-pressure tFUS applied to S1 in combination with vibration-tactile stimulation, using both high PRF, high DC, and low PRF, low DC protocols. High PRF, high DC, and high-pressure tFUS produced excitatory modulation of the sensory pathway, whereas low PRF, low DC, and low-pressure tFUS induced inhibitory effects. Mechanistically, these bidirectional effects were mediated primarily by S1 CaMKII-positive neurons, with excitatory modulation resulting from their activation and inhibitory modulation from their deactivation (Fig. 8). Overall, these results demonstrate that tFUS can bidirectionally modulate the cortico–thalamo–cortical sensory processing pathway by selectively engaging excitatory neurons, providing mechanistic insight into parameter-dependent neuromodulation and guiding the application of tFUS as a therapeutic tool for sensory disorders.

**Fig. 8.**
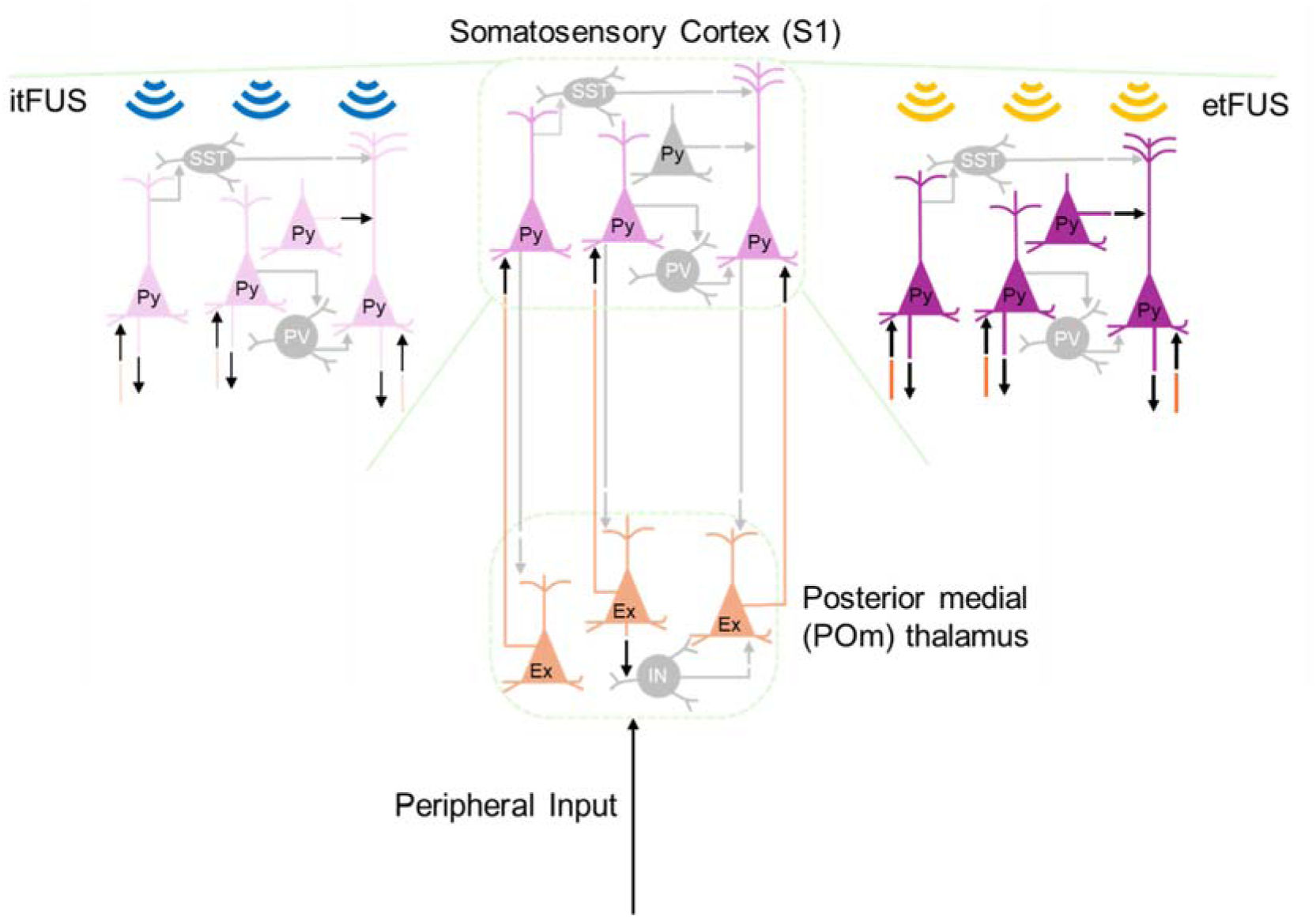
Summary of this work. Gray indicates resting-state activity. Peripheral input (vibration–tactile stimulation) excites CaMKII-positive neurons (Py) in S1 and excitatory neurons (Ex) in POm. Excitatory tFUS enhances sensory-related CTC pathway by activating CaMKII-positive neurons in S1 (indicated by darker colors and thicker arrows). Inhibitory tFUS suppresses this pathway by deactivating CaMKII-positive neurons in S1 (indicated by lighter colors and thinner arrows). Excitatory tFUS (etFUS) was delivered at a pulse repetition frequency (PRF) of 3 kHz with a 60% duty cycle at high pressure, whereas inhibitory tFUS (itFUS) was delivered at a PRF of 30 Hz with a 0.6% duty cycle at low pressure.

### Bidirectional tFUS parameter-dependent modulation

In this study, we demonstrated that tFUS exerts bidirectional, parameter-dependent modulatory effects on sensory processing. High PRF, high DC, and high-pressure stimulation produced excitatory effects, whereas low PRF, low DC, and low-pressure stimulation induced inhibitory effects (Fig. 4–5). The changes in sensory evoked potentials support tFUS-induced neural modulation. [16,27–29]. Legon et al. reported that tFUS applied to the S1 could attenuate amplitudes of somatosensory evoked potentials and enhance behavioral performance in sensory discrimination tasks[16]. Studies reported that the bidirectional neuromodulatory effects of tFUS depend critically on stimulation parameters, including PRF, DC, and acoustic pressure levels. For example, Yoon et al. demonstrated that high DC levels (30%, 50%, and 70%) tFUS targeting the motor cortex (M1) and thalamus induced excitatory effects, whereas low DC levels (3–5%) applied to S1 and the thalamus produced inhibitory responses [13]. In another study using different PRFs to stimulate the motor cortex of rodents, high PRF (>500 Hz) resulted in a marked increase in motor evoked potentials (MEP) [30]. In humans, Zadeh et al. reported that low PRF stimulation at 10 Hz produced inhibitory effects on MEP amplitudes[31]. Additionally, Kim et al. investigated the effects of tFUS on the mouse S1 using two different Ispta and observed increased hemodynamic response amplitudes with higher acoustic pressure[32]. However, increasing pressure levels may also raise safety concerns, such as potential neuronal damage or localized tissue heating [33]. Therefore, careful selection and optimization of tFUS parameters are essential to achieve the desired neuromodulatory effects while maintaining safety.

### Cortico-thalomo-cortical sensory processing pathway

Cortico-thalamo-cortical (CTC) pathways are highly involved in sensory processing, including visual, somatosensory, and auditory information [34]. These pathways involve precise interactions between specific cortical regions and corresponding thalamic nuclei, allowing for both feedforward transmission of ascending sensory signals and feedback modulation from cortex to thalamus [35]. In the somatosensory system, first-order thalamic nuclei, such as the ventral posterior medial nucleus (VPM), primarily relay peripheral sensory inputs to S1, whereas higher-order nuclei, including the posterior medial nucleus (POm), integrate inputs from multiple cortical and subcortical sources to mediate context-dependent processing, attention, and sensorimotor coordination [36–38]. Within these pathways, distinct neuron types contribute differentially: excitatory pyramidal neurons, such as CaMKII-positive neurons in S1, provide the principal output driving feedforward thalamocortical activation while in the thalamus, RSUs transmit excitatory signals back to cortex [38]. Consistent with this functional organization, our results showed that during vibration-tactile stimulation of the sensory pathway, only S1 CaMKII-positive neurons and POm RSUs responded robustly, whereas inhibitory interneurons (PV-positive and SST-positive) in S1 and FSUs in POm exhibited no responses (Figs. 3–6, Fig. S1). These findings highlight the critical role of excitatory neurons in mediating sensory information along the CTC pathway.

### Cell-type selectivity of tFUS

tFUS has been shown to exhibit cell-type selectivity in a parameter-dependent manner. Our results demonstrate that S1 CaMKII-positive neurons display distinct responses to tFUS under different stimulation parameters (Fig. 7, Fig. S2), consistent with previous studies showing that high PRF selectively excites excitatory neurons[17]. This selective responsiveness may arise from differences in membrane properties and ion channel composition across neuronal subtypes. Several studies have suggested that ultrasound neuromodulation interacts with mechanosensitive ion channels and voltage-gated channels embedded in neuronal membranes. For instance, low PRF tFUS has been reported to primarily affect potassium (K⁺) channels [39–42] whereas high PRF stimulation can additionally modulate sodium (Na⁺) and calcium (Ca²⁺) channels[43]. Because different neuronal populations express distinct combinations of these ion channels, they may exhibit differential sensitivity to specific tFUS parameters. Pyramidal neurons, including CaMKII-positive excitatory neurons in S1, express multiple tFUS-sensitive ion channels such as K⁺ [40]and Ca²⁺ [44]channels, which may account for their distinct responses to tFUS with different parameters.

### Future Applications

Bidirectional modulation of the sensory pathway could have therapeutic potential for various sensory-related disorders. For conditions characterized by overactive sensory processing, low-intensity tFUS could be used to suppress activity, such as sensory hypersensitivity in autism spectrum disorder; whereas high-intensity tFUS could enhance activity in cases of sensory deficits, such as stroke-induced sensory loss.

### Limitations and future work

In this study, only neuronal activation was assessed using optogenetics, which limits the precision of our conclusions regarding inhibitory effects. Future studies incorporating pharmacological approaches or optogenetic silencing of specific neuronal populations could provide stronger evidence for the mechanisms underlying tFUS-induced inhibition.

Only male subjects were used in this study. Although previous study has reported no sex differences in tFUS-induced pain modulation[11], sex-dependent differences in neuronal number and neuronal processes may still influence neuronal responses to tFUS and therefore need further investigation[45].

## Conclusions

By combining optogenetic tagging, multi-site electrophysiology, and waveform-based neuronal classification, we found that S1 CaMKII-positive neurons and POm RSUs responded robustly to vibration-tactile stimulation. We further demonstrated that tFUS applied to S1 can bidirectionally modulate the cortico–thalamo–cortical sensory pathway in a parameter-dependent manner: high PRF, high DC, and high-pressure stimulation elicited excitatory effects, whereas low PRF, low DC, and low-pressure stimulation produced inhibitory effects. Mechanistically, these bidirectional effects are primarily mediated by S1 CaMKII-positive neurons, with excitatory modulation driven by their activation and inhibitory modulation arising from their deactivation. This study provides mechanistic insight into the cell-type-specific effects of tFUS on sensory processing and highlights the critical role of CaMKII-positive neurons in mediating these effects. Importantly, these findings have translational implications, suggesting that tFUS parameters can be optimized to either enhance or suppress sensory processing, providing a foundation for the safe and effective development of tFUS-based therapies for sensory disorders.

## Funding

NIH NS124564, NS131069 (BH).

## Data availability

Experimental data supporting the findings will be made available in a public online repository when the paper is published.

## Acknowledgments

Some image components in the figures were created with BioRender.com.

## Author Contributions

**Huan Gao:** Writing - review & editing, Writing - original draft, Formal analysis, Data curation, Methodology, Investigation, Conceptualization. **Sandhya Ramachandran:** Writing - review & editing, Data curation, Investigation. **Mary M. Torregrossa:** Writing - review & editing, Methodology. **Bin He:** Writing - review & editing, Methodology, Investigation, Supervision, Conceptualization.

## Declaration of Competing Interest

The authors have no competing interests to declare.

## Notes

### Competing Interest Statement

The authors have declared no competing interest.

## References

[1] Gao H, Ramachandran S, Yu K, He B. Transcranial Focused Ultrasound Modulates Feedforward and Feedback Cortico-Thalamo-Cortical Pathways by Selectively Activating Excitatory Neurons. The Journal of Neuroscience 2025;45:e2218242025. 10.1523/JNEUROSCI.2218-24.2025.

[2] Mohammadjavadi M, Ye PP, Xia A, Brown J, Popelka G, Pauly KB. Elimination of peripheral auditory pathway activation does not affect motor responses from ultrasound neuromodulation. Brain Stimul 2019;12. 10.1016/j.brs.2019.03.005.

[3] Fine JM, Mysore AS, Fini ME, Tyler WJ, Santello M. Transcranial focused ultrasound to human rIFG improves response inhibition through modulation of the P300 onset latency. Elife 2023;12:e86190. 10.7554/eLife.86190.

[4] Kuhn T, Spivak NM, Dang BH, Becerra S, Halavi SE, Rotstein N, et al. Transcranial focused ultrasound selectively increases perfusion and modulates functional connectivity of deep brain regions in humans. Front Neural Circuits 2023;17. 10.3389/fncir.2023.1120410.

[5] Li X, Badran B, Dowdle L, Caulfield K, Summers P, Short B, et al. Imaged-guided Transcranial focused ultrasound on the right thalamus modulates ascending pain pathway to somatosensory cortex in healthy participants. Brain Stimul 2021;14. 10.1016/j.brs.2021.10.160.

[6] Yang H, Zhao Z, Xu R, Pei J, Yan J, Zhang X, et al. Low-Intensity Transcranial Ultrasound Stimulation Modulates the Excitability of Motor Cortical Neural Activity by Stimulating the Cerebellum. IEEE Transactions on Neural Systems and Rehabilitation Engineering 2025;33:3314–22. 10.1109/TNSRE.2025.3601110.

[7] Mishra A, Yang PF, Manuel TJ, Newton AT, Phipps MA, Luo H, et al. Disrupting nociceptive information processing flow through transcranial focused ultrasound neuromodulation of thalamic nuclei. Brain Stimul 2023;16. 10.1016/j.brs.2023.09.013.

[8] Bubrick EJ, McDannold NJ, Orozco J, Mariano TY, Rigolo L, Golby AJ, et al. Transcranial ultrasound neuromodulation for epilepsy: A pilot safety trial. Brain Stimul 2024;17. 10.1016/j.brs.2023.11.013.

[9] Succop Jr. BS, Seas A, Woo J, Bode Padron KJ, Bartlett AM, Shah B, et al. Focused Ultrasound in the Treatment of Epilepsy: Current Applications and Future Directions. Stereotact Funct Neurosurg 2025:166–88. 10.1159/000545716.

[10] Gao H, Frake A, Durand DM, He B. Transcranial Focused Ultrasound Stimulation Targeting White Matter Inhibits Seizures in a Rat Model of Epilepsy. IEEE Transactions on Neural Systems and Rehabilitation Engineering 2026;34. 10.1109/TNSRE.2025.3644273.

[11] Kim MG, Yu K, Yeh C-Y, Fouda R, Argueta D, Kiven S, et al. Low-intensity transcranial focused ultrasound suppresses pain by modulating pain-processing brain circuits. Blood 2024;144:1101–15. 10.1182/blood.2023023718.

[12] Pang N, Wang Q, Zhang H, Su R, Yan J, Yuan Y. Effects of Low-Intensity Transcranial Ultrasound Stimulation on Cortical Functional Network Connections and Epileptic Seizures in a Mouse Model of Epilepsy. IEEE Transactions on Neural Systems and Rehabilitation Engineering 2026;34:1176–86. 10.1109/TNSRE.2026.3664698.

[13] Yoon K, Lee W, Lee JE, Xu L, Croce P, Foley L, et al. Effects of sonication parameters on transcranial focused ultrasound brain stimulation in an ovine model. PLoS One 2019;14. 10.1371/journal.pone.0224311.

[14] Dallapiazza RF, Timbie KF, Holmberg S, Gatesman J, Lopes MB, Price RJ, et al. Noninvasive neuromodulation and thalamic mapping with low-intensity focused ultrasound. J Neurosurg 2018;128. 10.3171/2016.11.JNS16976.

[15] Liu C, Yu K, Niu X, He B. Transcranial Focused Ultrasound Enhances Sensory Discrimination Capability through Somatosensory Cortical Excitation. Ultrasound Med Biol 2021;47. 10.1016/j.ultrasmedbio.2021.01.025.

[16] Legon W, Sato TF, Opitz A, Mueller J, Barbour A, Williams A, et al. Transcranial focused ultrasound modulates the activity of primary somatosensory cortex in humans. Nat Neurosci 2014;17:322–9. 10.1038/nn.3620.

[17] Yu K, Niu X, Krook-Magnuson E, He B. Intrinsic functional neuron-type selectivity of transcranial focused ultrasound neuromodulation. Nat Commun 2021;12:2519. 10.1038/s41467-021-22743-7.

[18] Lemaire T, Yuan Y, Gellman C, LeMessurier AM, Haiken Dray SR, Little JP, et al. Microscopic deconstruction of cortical circuit stimulation by transcranial ultrasound. BioRxiv 2024:2024.10.10.617091. 10.1101/2024.10.10.617091.

[19] Paxinos G, Watson C. The rat brain in stereotaxic coordinates: hard cover edition. Elsevier; 2006.

[20] Mo C, Sherman SM. A sensorimotor pathway via higher-order thalamus. Journal of Neuroscience 2019;39:692–704. 10.1523/JNEUROSCI.1467-18.2018.

[21] Lee KS, Loutit AJ, de Thomas Wagner D, Sanders M, Prsa M, Huber D. Transformation of neural coding for vibrotactile stimuli along the ascending somatosensory pathway. Neuron 2024;112. 10.1016/j.neuron.2024.07.005.

[22] Callier T, Gitchell T, Harvey MA, Bensmaia SJ. Disentangling Temporal and Rate Codes in the Primate Somatosensory Cortex. Journal of Neuroscience 2024;44. 10.1523/JNEUROSCI.0036-24.2024.

[23] Bieler M, Xu X, Marquardt A, Hanganu-Opatz IL. Multisensory integration in rodent tactile but not visual thalamus. Sci Rep 2018;8. 10.1038/s41598-018-33815-y.

[24] Lakunina A, Socha KZ, Ladd A, Bowen AJ, Chen S, Colonell J, et al. Neuropixels Opto: Combining high-resolution electrophysiology and optogenetics. BioRxiv 2025. https://www.biorxiv.org/10.1101/2025.02.04.636286v3

[25] Chung S, Weber F, Zhong P, Tan CL, Nguyen TN, Beier KT, et al. Identification of preoptic sleep neurons using retrograde labelling and gene profiling. Nature 2017;545. 10.1038/nature22350.

[26] Kim MG, Yu K, Niu X, He B. Investigation of Displacement of Intracranial Electrode Induced by Focused Ultrasound Stimulation. IEEE Trans Instrum Meas 2021;70. 10.1109/TIM.2021.3125978.

[27] Lee W, Kim H, Jung Y, Song I-U, Chung YA, Yoo S-S. Image-Guided Transcranial Focused Ultrasound Stimulates Human Primary Somatosensory Cortex. Sci Rep 2015;5:8743. 10.1038/srep08743.

[28] Kim HC, Lee W, Weisholtz DS, Yoo SS. Transcranial focused ultrasound stimulation of cortical and thalamic somatosensory areas in human. PLoS One 2023;18. 10.1371/journal.pone.0288654.

[29] Darrow DP, O’Brien P, Richner TJ, Netoff TI, Ebbini ES. Reversible neuroinhibition by focused ultrasound is mediated by a thermal mechanism. Brain Stimul 2019;12. 10.1016/j.brs.2019.07.015.

[30] King RL, Brown JR, Newsome WT, Pauly KB. Effective parameters for ultrasound-induced in vivo neurostimulation. Ultrasound Med Biol 2013;39. 10.1016/j.ultrasmedbio.2012.09.009.

[31] Zadeh AK, Raghuram H, Shrestha S, Kibreab M, Kathol I, Martino D, et al. The effect of transcranial ultrasound pulse repetition frequency on sustained inhibition in the human primary motor cortex: A double-blind, sham-controlled study. Brain Stimul 2024;17:476–84. 10.1016/j.brs.2024.04.005.

[32] Kim E, Anguluan E, Youn S, Kim J, Hwang JY, Kim JG. Non-invasive measurement of hemodynamic change during 8 MHz transcranial focused ultrasound stimulation using near-infrared spectroscopy. BMC Neurosci 2019;20. 10.1186/s12868-019-0493-9.

[33] Nandi T, Kop BR, Butts Pauly K, Stagg CJ, Verhagen L. The relationship between parameters and effects in transcranial ultrasonic stimulation. Brain Stimul 2024;17. 10.1016/j.brs.2024.10.008.

[34] Guillamón-Vivancos T, Aníbal-Martínez M, Puche-Aroca L, Martini FJ, López-Bendito G. Sensory modality-specific wiring of thalamocortical circuits. Nat Rev Neurosci 2025;26. 10.1038/s41583-025-00945-y.

[35] Groh A, Mease R. Corticothalamic Pathways in the Somatosensory System. The Thalamus, Cambridge University Press; 2022, p. 221–36. 10.1017/9781108674287.013.

[36] Sanganahalli BG, Herman P, Rothman DL, Blumenfeld H, Hyder F. Metabolic demands of neural-hemodynamic associated and disassociated areas in brain. Journal of Cerebral Blood Flow and Metabolism 2016;36. 10.1177/0271678X16664531.

[37] Pais-Vieira M, Lebedev MA, Wiest MC, Nicolelis MAL. Simultaneous top-down modulation of the primary somatosensory cortex and thalamic nuclei during active tactile discrimination. Journal of Neuroscience 2013;33. 10.1523/JNEUROSCI.1659-12.2013.

[38] O’Reilly C, Iavarone E, Yi J, Hill SL. Rodent somatosensory thalamocortical circuitry: Neurons, synapses, and connectivity. Neurosci Biobehav Rev 2021;126. 10.1016/j.neubiorev.2021.03.015.

[39] Kubanek J, Shi J, Marsh J, Chen D, Deng C, Cui J. Ultrasound modulates ion channel currents. Sci Rep 2016;6:24170. 10.1038/srep24170.

[40] Lin Z, Huang X, Zhou W, Zhang W, Liu Y, Bian T, et al. Ultrasound Stimulation Modulates Voltage-Gated Potassium Currents Associated With Action Potential Shape in Hippocampal CA1 Pyramidal Neurons. Front Pharmacol 2019;10. 10.3389/fphar.2019.00544.

[41] Sorum B, Rietmeijer RA, Gopakumar K, Adesnik H, Brohawn SG. Ultrasound activates mechanosensitive TRAAK K+ channels through the lipid membrane. Proceedings of the National Academy of Sciences 2021;118:e2006980118. 10.1073/pnas.2006980118.

[42] Clennell B, Steward TGJ, Hanman K, Needham T, Benachour J, Jepson M, et al. Ultrasound modulates neuronal potassium currents via ionotropic glutamate receptors. Brain Stimul 2023;16. 10.1016/j.brs.2023.01.1674.

[43] Morris CE, Juranka PF. Nav channel mechanosensitivity: Activation and inactivation accelerate reversibly with stretch. Biophys J 2007;93. 10.1529/biophysj.106.101246.

[44] Almog M, Korngreen A. Characterization of Voltage-Gated Ca2+ Conductances in Layer 5 Neocortical Pyramidal Neurons from Rats. PLoS One 2009;4:e4841-. 10.1371/journal.pone.0004841

[45] Rabinowicz T, Dean DE, Petetot JMDC, De Courten-Myers GM. Gender differences in the human cerebral cortex: More neurons in males; more processes in females. J Child Neurol 1999;14. 10.1177/088307389901400207.

