## Supplementary Figures for "Cell-Type-Specific Bidirectional Modulation of the Cortico–Thalamo–Cortical Sensory Pathway by Transcranial Focused Ultrasound (tFUS)"

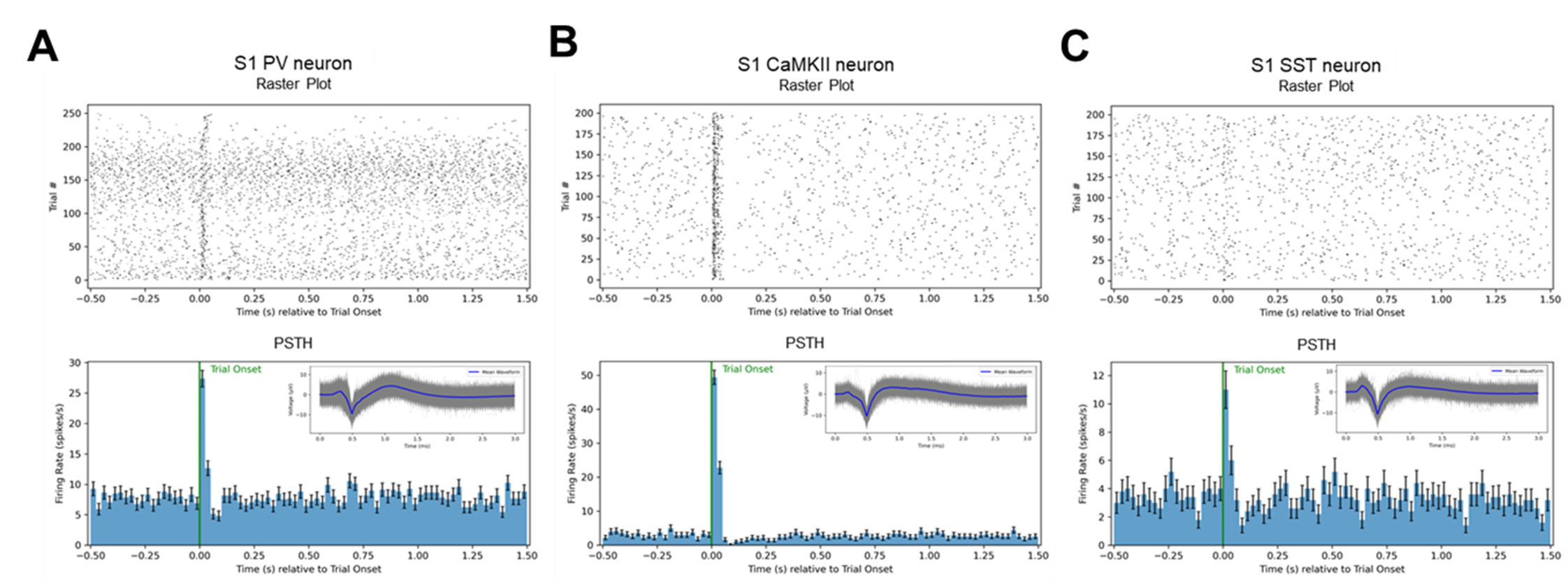

**Fig. S1.** Representative responses of S1 PV-positive (**A**), CaMKII-positive (**B**), and SST-positive (**C**) neurons to optical stimulation for neuronal identification. Upper panels show raster plots across trials; lower panels show peri-event spike rate histograms, with inset panels displaying the averaged spike waveform.

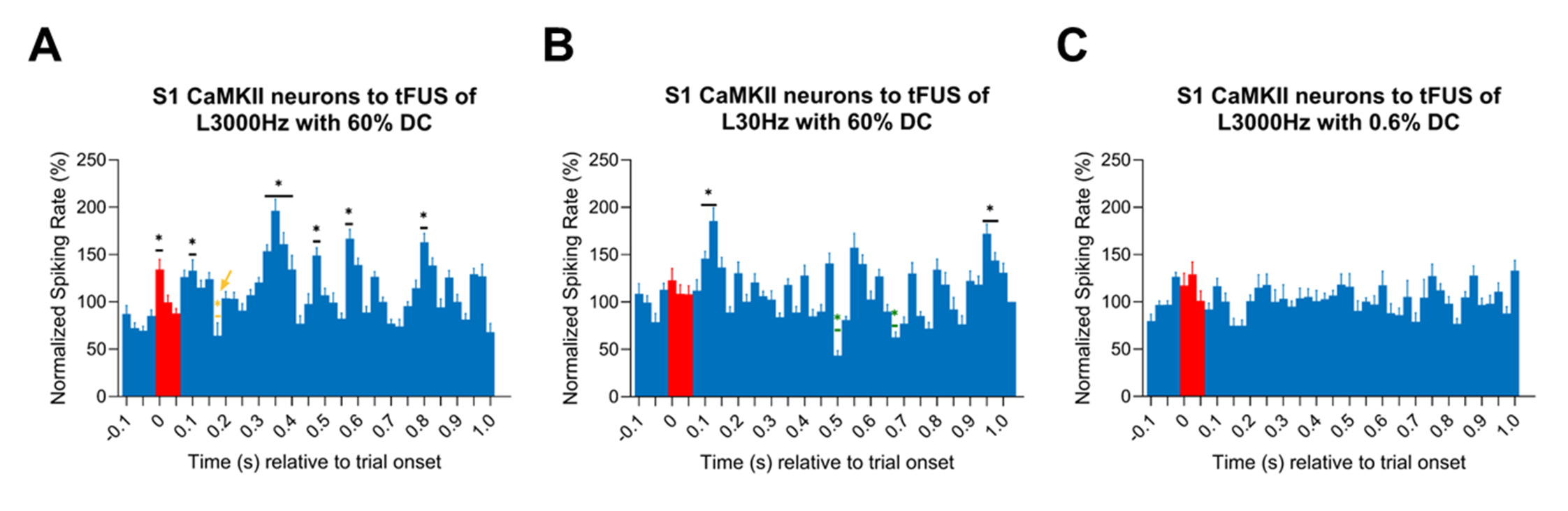

**Fig. S2.** S1 CaMKII-positive neuronal responses to low-pressure tFUS under different stimulation parameters. (A) 3000 Hz PRF with 60% duty cycle, (B) 30 Hz PRF with 60% duty cycle, and (C) 3000 Hz PRF with 0.6% duty cycle. Yellow * (pointed by yellow arrow) indicates a significant decrease in normalized firing rate compared to the 100 ms pre-stimulation baseline, whereas black * indicates significant increases relative to baseline (p < 0.05). Bin size was 25 ms. Red bars denote bin windows during the stimulation period.
